# Anterior temporal lobe morphometry predicts categorization ability

**DOI:** 10.1101/214965

**Authors:** B. Garcin, M. Urbanski, M. Thiebaut De Schotten, R. Levy, E. Volle

## Abstract

Categorization is the mental operation by which the brain classifies objects and events. It is classically assessed using semantic and non-semantic matching or sorting tasks. These tasks show a high variability in performance across healthy controls and the cerebral bases supporting this variability remain unknown. In this study we performed a voxel-based morphometry study to explore the relationships between semantic and shape categorization tasks and brain morphometric differences in 50 controls. We found significant correlation between categorization performance and the volume of the grey matter in the right anterior middle and inferior temporal gyri. Semantic categorization tasks were associated with more rostral temporal regions than shape categorization tasks. A significant relationship was also shown between white matter volume in the right temporal lobe and performance in the semantic tasks. Tractography revealed that this white matter region involved several projection and association fibers, including the arcuate fasciculus, inferior fronto-occipital fasciculus, uncinate fasciculus, and inferior longitudinal fasciculus. These results suggest that categorization abilities are supported by the anterior portion of the right temporal lobe and its interaction with other areas.

**Highlights:**

- Anterior temporal lobe morphometry correlates with categorization performances
- Semantic is associated with a more rostral temporal region than shape categorization
- Semantic categorization performances are associated with right temporal connections

## 1. Introduction

Categorization is the mental operation by which the brain classifies objects and events. The ability to categorize information has an impact in virtually all domains of cognition and behavior, from learning (children learn new concepts by categorizing items that look similar or have similar properties) to survival (to recognize an animal as dangerous, primates need to categorize it as similar to a previously encountered dangerous animal).

The evaluation of categorization abilities relies on various tests, including semantic and visual categorization tests. Semantic categorization abilities are usually assessed by matching tests based on taxonomic or thematic categorization, such as the Pyramid and Palm Tree Test (PPT test) (Howard and Patterson 1992), and by the production of the relevant abstract category as in the similarities subtest of the Wechsler Intelligence Adult Scale (WAIS) (Wechsler 2008). Categorization abilities can also be assessed by sorting tests such as the Delis–Kaplan Executive Function (D-KEFS) sorting test (Delis et al. 2001) and the Wisconsin Card Sorting Test (WCST) (Milner 1963), that use semantic and/or visuoperceptual material. All of these tests but the WAIS are designed to screen and detect deficits in patients. However, normative studies focused on these tests show inter-individual variability in healthy subjects’ performances with a relative standard deviation (i.e. rSD = standard deviation/mean) of 4%–12% in the PPT test (Howard and Patterson 1992; Rami et al. 2008; Klein and Buchanan 2009; Gamboz et al. 2009; Callahan et al. 2010), an rSD of 25%–45% in the similarities subtest of the WAIS (Wechsler 2008; Wisdom et al. 2012; Harrison et al. 2014), an rSD of 20%–40% in the D-KEFS sorting test (Delis et al. 2001; Homack et al. 2005; Mattioli et al. 2014), and an rSD of 20%–60% in the number of categories found in the WCST (Caffarra et al. 2004; Shan et al. 2008; Arango-Lasprilla et al. 2015). Whether this inter-individual variability in categorization tasks is related to variability in brain structure remains unknown.

Functional neuroimaging studies in healthy subjects, as well as electrophysiological studies in primates, have shown the involvement of various brain regions in categorization tasks. For instance, the ventrolateral prefrontal cortex (PFC) (Vogels et al. 2002; Grossman et al. 2002; Koenig et al. 2005; Milton et al. 2009), the lateral and/or inferior temporal cortices (Gerlach et al. 2000; Sigala and Logothetis 2002; Pernet et al. 2005), or both frontal and temporal cortices (Tyler et al. 2001; Devlin et al. 2002; Adams and Janata 2002; Pilgrim et al. 2002; Reber et al. 2002; Pernet et al. 2004; Sass et al. 2009; Visser et al. 2012) are involved during semantic and visuoperceptual categorization tasks. Some authors used distinct task instructions to explore executive control processes separately from bottom-up access to visuoperceptual and semantic representations (Koenig et al. 2005; Milton et al. 2009; Garcin et al. 2012). For instance, Garcin et al used matching and non-matching sorting tasks and showed that BOLD signal was higher in the ventrolateral PFC for the matching than the non-matching tasks suggesting that matching involves more control processes than non-matching (Garcin et al. 2012). All these studies examined the regions activated during categorization, without assessing the relationship between brain structural variability and categorization abilities.

Regarding brain structure, the exact shape of every human brain is unique, resulting in inter-individual anatomical variability (Mazziotta et al. 1995; Uylings et al. 2005; Fischl et al. 2008), but whether inter-individual variability can affect or predict individual categorization performance is unknown. We hypothesized that structural variations in the regions classically observed in functional imaging (the lateral prefrontal cortex and the lateral and inferior temporal cortices) may be related to subjects’ performance in categorization. To address this question, we performed a voxel-based morphometry (VBM) study in healthy subjects using a sorting test adapted from the PPT test (Howard and Patterson 1992) that allowed us to assess separately semantic and visuoperceptual categorization in matching and non-matching conditions.

## 2. Materials and methods

### 2.1. Participants

Fifty right-handed native French speakers (25 females; age 22–71 years, mean = 47±14.3 years) participated in the study. A large age range was chosen to represent the diversity of the general population. All participants were healthy adults with no history of neurological or psychiatric disorders and no abnormalities were revealed on their structural MRI. Participants had an average of 15.4±3.0 years of education (range, 10–26). They had no cognitive impairment as assessed with the Mini Mental State Examination (Folstein et al. 1975) and the Frontal Assessment Battery (Dubois et al. 2000). They all underwent a French verbal semantic matching test adapted from the word-written version of the PPT test (Merck et al. 2011) and showed no impairment. The experiment was approved by the local ethics committee. All participants provided written informed consent and were paid for their participation.

### 2.2. Experimental stimuli, tasks, and procedure

We used a short version of the categorization paradigm described in a previous functional imaging study (Garcin et al. 2012). The principle of this task is similar to that of the PPT test, a semantic matching task designed to search for semantic deficits in patients. Compared to the PPT, the categorization paradigm was designed to assess both semantic and visuoperceptual categorization, with two distinct sorting conditions: matching and non-matching conditions. The paradigm used a factorial design with two dimensions (i.e., *Shape and Category*) assessing semantic (*Category*) and visuoperceptual (*Shape*) categorization, and two conditions (*i.e., Same and Different*) assessing matching (*Same*) and non-matching (*Different*) sorting.

#### 2.3.1. Stimuli

Stimuli consisted of triads of black-and-white drawings of real-life objects that were displayed on a computer screen. One drawing at the top of the screen was framed; the two other drawings were located at the bottom left and right sides of the screen (Figure 1). For each trial, there was a semantic link between the framed drawing and one of the two bottom ones, as well as a similarity of shape between the framed drawing and one of the two bottom ones (for more information, see the legend of Figure 1). Of the 576 stimuli used in our previous fMRI study (Garcin et al. 2012), 160 stimuli were selected to create a shorter version of the paradigm. Stimuli belonged to 107 different categories, among which 60% were taxonomic (e.g., fruits or insects, *n* = 64), and 40% were thematic (e.g., rugby or transportation, *n* = 43). Among all drawings, 60% were non-living objects and 40% were living objects. Some objects were easy to handle (e.g., tools, fruit), and others were not (e.g., buildings, wild animals). See Supplementary material 1.

**Fig. 1.**
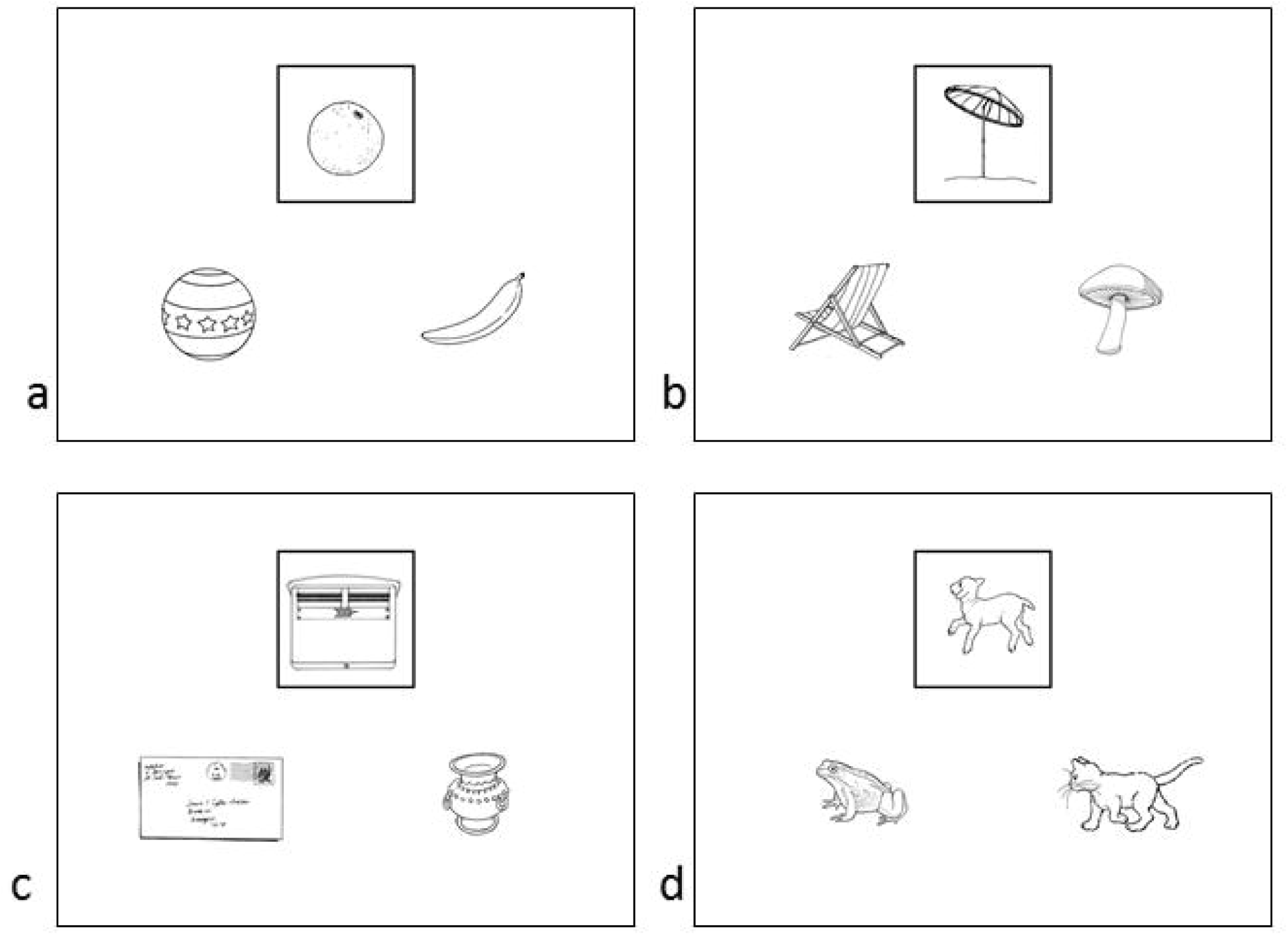
Samples of stimuli. The framed drawing was compared with the two bottom ones according to four possible instructions: *Same Shape, Same Category, Different Shape,* and *Different Category*. There was systematically an abstract and/or a shape relationship between the framed drawing and at least one of the two others. In half of the stimuli, one drawing had a similar shape, whereas the other one belonged to the same category as the framed drawing, such as in stimuli a and b. In stimulus a, the bottom right drawing belonged to the same category as the framed drawing (“fruits”), and the bottom left drawing was of the same shape (“round”). In stimulus b, the bottom right drawing was of the same shape as the framed drawing, and the bottom left belonged to the same category. In the other half, the drawing with the most similar shape belonged to the same category as the framed one, such as in stimuli c and d. Some categories (60%) were taxonomic, such as in stimuli 1 (“fruits”) and d (“mammals”), while others (40%) were thematic, such as in stimuli b and c (“contextual and functional link”).

#### 2.2.2. Experimental task

The 160 stimuli were divided into four sets of 40 stimuli each. Each set was assigned to one of the four following tasks: the *same shape* task, *same category* task, *different shape* task, and *different category* task. In the *same shape* task, participants had to choose the drawing that had the most similar shape to that of the framed drawing. In the *same category* task, participants had to choose the drawing that belonged to the same category as the framed drawing. In the *different shape* task, participants were asked to choose the drawing that had the most different shape from that of the framed drawing. In the *different category* task, participants had to choose the drawing that did not belong to the same category as the framed drawing. Correct items were equally distributed between the bottom-left and bottom-right drawings for each task. In each task, one bottom drawing had both the same shape and the same category as the framed drawing in half of the trials, whereas one bottom drawing had a similar shape and the other one belonged to the same category in the other half of the trials. This step ensured that shape had no effect on category decision and vice versa. The number of categories and their nature (taxonomic/thematic, living/non-living) were equally distributed in the four tasks.

#### 2.2.3. Experimental procedure

Stimulus presentation was programmed on a PC using meyeParadigm 1.17 software (<u>www.eye-brain.com</u>). The order of the tasks and the order of trials within each task were randomized between subjects, and each task was performed in a block of 40 trials. Training was performed before the beginning of the test. The instruction was given orally before each block, and it was reminded on the screen during 5 s at the beginning of each block. Participants had a maximum time of 10 s to answer to each stimulus, and a reminder of the instruction appeared during 1.5 s between each trial. Subjects had to press the E key to choose the left drawing and O key for the right drawing. Participants were asked to answer as fast and as accurate as possible. The total duration of the procedure was between 20 and 25 min. A quick debriefing was performed after each block.

### 2.3. Behavioral analysis

Accuracy and response times (RTs) were measured and statistical analyses were conducted using SPSS software (<u>http://www-01.ibm.com/software/analytics/spss/</u>). Repeated measures two-way ANOVA analyses were performed to compare participants’ performance (accuracy and RTs for correct responses only) according to two factors: dimension (*Category/Shape*) and condition *(Same/Different*). We also ran Pearson correlation analyses between age and experimental scores, as well as between education and experimental scores. We compared the performance of men and women using an independent samples *t*-test.

### 2.4. VBM study: Image acquisition and analysis

#### 2.4.1. Structural T1-weighted images

All participants underwent the same high-resolution T1-weighted structural MRI scans acquired on a Siemens 3 Tesla VERIO TIM system equipped with a 32-channel head coil. An axial 3D MPRAGE dataset covering the whole head was acquired for each participant as follows: 176 slices, voxel resolution = 1 mm × 1 mm × 1 mm, TE = 2.98 ms, TR = 2300 ms, flip angle = 9°.

#### 2.4.2. VBM pre-processing

3D T1-weighted sequences were processed and analyzed with SPM8 (Wellcome Department of Imaging Neuroscience, London, UK) running on Matlab (Mathworks Inc., USA; www.mathworks.com/matlabcentral). We used the VBM8 toolbox (http://dbm.neuro.uni-jena.de/vbm/) to perform MRI data pre-processing (http://dbm.neuro.uni-jena.de/vbm8/BVM8-Manual.pdf). First, we spatially normalized the T1 images to the MNI152 Dartel template using high-dimensional Dartel normalization (Ashburner 2007). SPM8’s new version of the unified segmentation method (new segment) (Ashburner and Friston 2005) was used to segment T1 images into grey matter (GM), white matter (WM), and cerebrospinal fluid. Default estimation parameters were used to compute normalized images with an isotropic voxel size of 1.5 mm^3^. Normalized images were modulated to compensate for regional volume changes caused by normalization. The “normalized non-linear modulation only” option was used, allowing us to analyze relative differences in regional GM volume corrected for individual brain size. The quality was evaluated by displaying one slice for each image module and searching for visual abnormalities and by checking sample homogeneity using the covariance between individual images. The images with low covariance (−2 standard deviations, *n* = 4) were visually examined, and none of them had to be excluded. In addition, all normalized 3D images were visually inspected and compared with the template using frontal anatomical landmarks by an expert neurologist (B.G.). Modulated and normalized GM images were then smoothed using a Gaussian kernel of 8 mm^3^ full width at half maximum to enable interindividual comparisons and parametric statistics. The resulting GM images were used for statistical analyses.

#### 2.4.3. VBM whole-brain statistical analysis

To investigate the relationship between VBM regional grey matter (GM) structural variability and different aspects of categorization, we ran multiple regression analyses in SPM8 between GM volume and behavioral scores. RTs for accurate responses were used for the analyses because of a ceiling effect in accuracy. First, the averaged scores in the Category dimension (*same category* and *different category* tasks) and the averaged scores in the Shape dimension (*same shape* and *different shape* tasks) were entered separately as covariates in two separate regression models. In a second step, the averaged scores in the Same conditions (*same category* and s*ame shape* tasks) and the averaged scores in the Different conditions (*different category* and *different shape* tasks) were entered separately as covariates in separate regression models. Age, gender, and education were co-varied out in all the regression models. Data were also normalized and corrected for individual total GM volume by entering their global values as covariates in the linear model. Global values of total GM volume were extracted and calculated from the get_totals scripts (available <u>http://www0.cs.ucl.ac.uk/staff/g.ridgway/vbm/get_totals.m</u>). For each regression analysis, we investigated significant results at p < 0.05 using a familywise error (FWE) correction at the cluster level with a voxel-level threshold of p < 0.001 uncorrected. Non-stationary smoothness of the data was taken into account for cluster-level threshold. Results at p < 0.001 uncorrected for multiple comparisons at the voxel level, with a minimal cluster size of 100 voxels, are reported in the supplementary results for information purposes.

To investigate the relationship between VBM regional white matter (WM) density and different aspects of categorization, we ran multiple regression analyses in SPM8 between WM volume and behavioral scores. We used the same models and covariates as for the GM VBM analyses. Data were also normalized and corrected for individual total WM volume by entering their values as covariates in the linear model. For each regression analysis, we investigated significant results at p < 0.05 using FWE correction at the cluster level with a voxel-level threshold of p < 0.001 uncorrected. Non-stationary smoothness of the data was taken into account for cluster-level threshold. Results at *p* < 0.001 uncorrected for multiple comparisons at the voxel level, with a minimal cluster size of 100 voxels are reported in the supplementary results.

### 2.5 Connectivity study: image acquisition, preprocessing, and analysis

The functions of brain regions depend on their connectivity with other brain regions. Therefore, anatomical connectivity of the VBM results was investigated in a connectivity study using diffusion images. We explored the connections terminating in and emerging from the brain regions identified in the WM VBM in 44 out of the 50 participants (22 females; age 22–71 years, mean = 46.5±14.5 years).

#### 2.5.1 Diffusion image acquisition

A total of 70 near-axial slices were acquired during the same MRI session as T1 images. We used an acquisition sequence fully optimized for tractography of DWI that provided isotropic (2 mm × 2 mm × 2 mm) resolution and coverage of the whole head. The acquisition was peripherally gated to the cardiac cycle with an echo time (TE) of 85 ms. We used a repetition time (TR) equivalent to 24 RR. At each slice location, six images were acquired with no diffusion gradient applied. Sixty diffusion-weighted images were acquired in which gradient directions were uniformly distributed in space. Diffusion weighting was equal to a b-value of 1500 s/mm^2^.

#### 2.5.2 Diffusion imaging pre-processing

One supplementary image with no diffusion gradient applied but with reversed phase-encode blips was collected. This step provided us with a pair of images with no diffusion gradient applied and distortions going in opposite directions. From these pairs, the susceptibility-induced off-resonance field was estimated using a method similar to that described in (Andersson et al. 2003) and corrected on the whole diffusion-weighted dataset using the tool TOPUP as implemented in FSL (Smith et al. 2004). Finally, at each slice, diffusion-weighted data were simultaneously registered and corrected for subject motion and geometrical distortion, adjusting the gradient accordingly (ExploreDTI http://www.exploredti.com) (Leemans and Jones 2009).

#### 2.5.3 Spherical deconvolution tractography reconstruction

Spherical deconvolution was chosen to estimate multiple orientations in voxels containing different populations of crossing fibers (Tournier et al. 2004; Anderson 2005). The damped version of the Richardson–Lucy algorithm for spherical deconvolution (Dell’acqua et al. 2010) was calculated using an in-house developed software. Algorithm parameters were chosen as previously described (Dell’acqua et al. 2012).

Whole-brain tractography was performed by selecting every brain voxel with at least one fiber orientation as a seed voxel. From these voxels and for each fiber, orientation streamlines were propagated using Euler integration with a step size of 1 mm. When entering a region with crossing WM bundles, the algorithm followed the orientation vector of the least curvature (Schmahmann et al. 2007). Streamlines were halted when a voxel without fiber orientation was reached or when the curvature between two steps exceeded a threshold of 60°. Spherical deconvolution, fiber orientation vector estimation, and tractography were performed using in-house software developed with Matlab 7.8 (http://www.mathworks.com).

#### 2.5.4 Tractography dissections

The significant results of WM VBM analysis were used as regions of interest (ROIs) for tract dissections. We dissected the tracts connecting the observed ROIs associated with *Category* (i.e., *same category* + *different category*) performances.

In short, each participant’s convergence speed maps (Dell’acqua et al. 2012) were registered to the MNI152 template using Advanced Normalization Tools (Klein et al. 2009). Inverse deformation was then applied to the ROIs to bring them within the native space of each participant. Binary individual visitation maps were created for the connections emerging from or terminating in the observed ROI by assigning each voxel a value of 1 or 0, depending on whether the voxel was intersected by the streamlines of the tract. Binary visitation maps of each of the dissected tracts were normalized to the MNI space using the same affine and diffeomorphic deformations as calculated above. We created percentage overlap maps by adding the normalized visitation maps from each subject at each point in the MNI space. Therefore, the overlap of the visitation maps varies according to inter-subject variability. We inspected tracts reproducible in more than 50% of the participants using a method described previously in (Thiebaut de Schotten et al. 2011). Tracts resulting from this analysis were visually inspected and identified using an atlas of human brain connections (Thiebaut de Schotten et al. 2011; Rojkova et al. 2015).

## 3 Results

### 3.1. Behavioral Results

#### 3.1.1. Accuracy (Figure 2a)

The mean error rate was low (mean: 3.2%, all conditions included). Repeated measures two-way ANOVAs revealed no effect of dimension (i.e., *Category* vs. *Shape*; F(1,49) = 0.98, p = 0.32) or condition (i.e., *Same* vs. *Different*; F(1,49) = 0.47, p = 0.49).

**Fig. 2.**
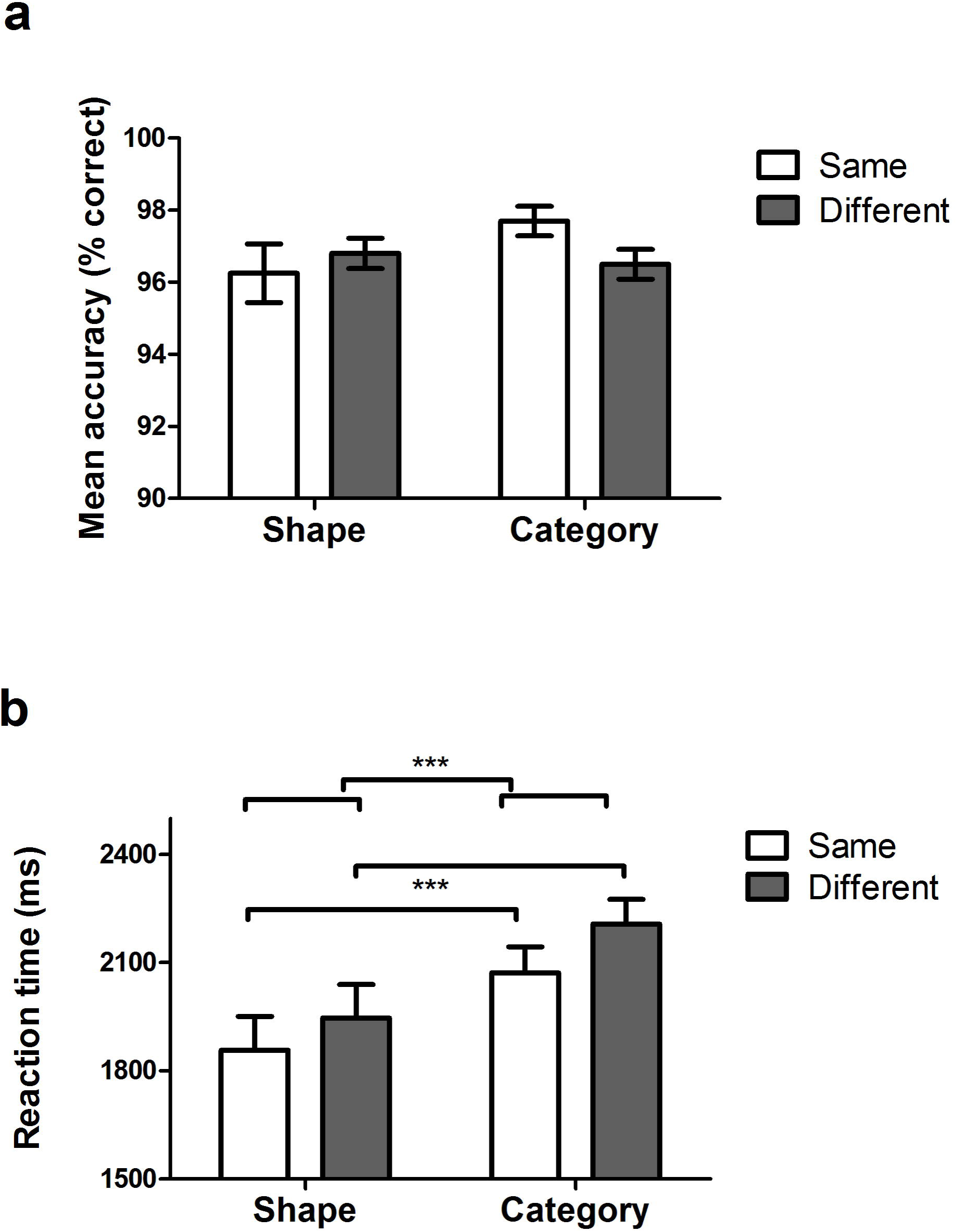
Behavioral data. Histograms represent means ± standard errors of the mean. ***: p ≤ 0.001. a. Accuracy in *Shape*, *Category, Same*, and *Different* tasks. Repeated measures two-way ANOVAs revealed no effect of dimension (i.e., Category vs. Shape) or condition (i.e., Same vs. Different). b. RTs for S*hape*, *Category, Same*, and *Different* tasks. Repeated measures two-way ANOVA revealed a significant effect of dimension (Shape vs. Category tasks, p < 0.001) and a significant effect of condition (Same vs. Different tasks, p = 0.001). No significant interaction was found between dimension and condition.

#### 3.1.2. RTs (Figure 2b)

Repeated measures two-way ANOVA revealed a significant effect of dimension (F(1,49) = 18.7, p < 0.001, *Shape* mean = 1902 ms, *Category* mean = 2140 ms) and a significant effect of condition (F(1,49) = 12.7, p = 0.001, *Same* mean = 1965 ms, *Different* mean = 2077 ms). No significant interaction was found between dimension and condition (F(1, 49) = 0.39, p = 0.53).

#### 3.1.3. Correlations: Age, Gender, and Education

Age was significantly positively correlated with RT in all conditions: *same shape* (r = 0.50, p < 0.001), *different shape* (r = 0.55, p <0.001), *same category* (r = 0.47, p = 0.001), and *different category* (r = 0.47, p = 0.001). There was no significant gender difference for each task. Education was not correlated with RT in any tasks or with average RT of all tasks pooled together.

### 3.2. GM correlations with RT in the Shape and Category dimensions (Table 1, Figure 3, supplementary figure 1)

Voxel-wise multiple regression analyses of RTs for each task dimension (*Shape* and *Category*) were conducted within GM with age, gender, and education as covariates of non-interest.

**Table 1.** VBM–whole brain analysis: negative GM correlations with RT in Shape, Category, Same, and Different tasks at p < 0.05 after FWE correction at the cluster level.

|  | brain region | Side | BA | MNI coordinate<br>(maxima) | T value | cluster<br>size | cluster-level<br>p(FWE) |
| --- | --- | --- | --- | --- | --- | --- | --- |
| Shape | middle and inferior temporal gyrus | R | 20/21 | 56 -19 -20 | 4.74 | 679 | 0.044 |
| Category | Temporal pole, middle and inferior<br>temporal gyrus | R | 20/21/38 | 57 -2 -27 | 4.97 | 1558 | 0.003 |
| Same | temporal pole, middle and inferior<br>temporal gyrus | R | 20/21 | 57 -13 -20 | 5.00 | 1352 | 0.004 |
| Different | middle and inferior temporal gyrus | R | 20/21 | 57 -16 -21 | 4.41 | 1308 | 0.009 |

**Fig. 3.**
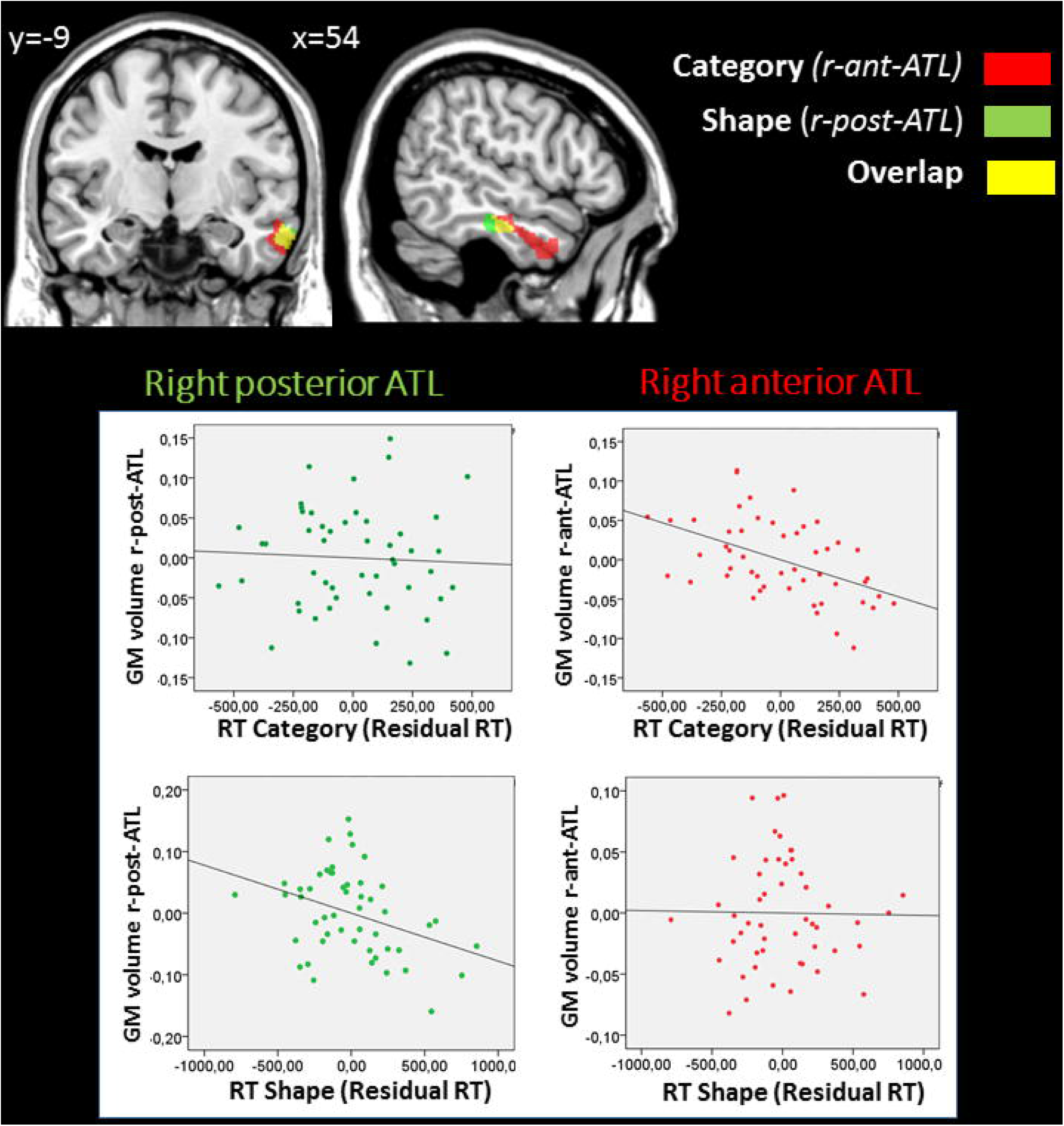
Results from the whole-brain GM VBM analysis according to dimension. p<0.05 after FWE correction. Significant regions associated with changes in GM volume related to performance in terms of RT are superimposed on a coronal (left) and sagittal (right) view. Additional slices can be found in supplementary figure 1. The whole-brain analyses identified a right anterior temporal region (r-ant-ATL) (in red), in which GM volume was negatively correlated with RT in the Category dimensions (*same and different category* tasks) and a most posterior ATL region (r-post-ATL) (in green) in which GM volume was negatively correlated with RT in the Shape (*same and different shape* tasks) dimensions. Shared regions are shown in yellow. Plots between performance and GM measures within these 2 regions are displayed in the partial regression diagrams: X axes represent the residual RT in each experimental dimension, and Y-axes the residual of the mean GM volume within each region. This analysis showed that the r-ant-ATL is significantly associated with *Category* but not *Shape*, while the r-post-ATL is significantly associated with *Shape* but not *Category*.

At a FWE-corrected threshold, RTs in the *Shape* and *Category* dimensions were both negatively correlated with GM volume in the right temporal lobe, i.e., less GM volume was related to slower RTs. RTs in the *Category* dimension were correlated with the right temporal pole, middle temporal gyrus, and inferior temporal gyrus (BA 20/21/38). RTs in the *Shape* dimension were correlated with the right middle temporal and inferior temporal gyri (BA 20/21). As Figure 3 shows, RTs in the *Category* dimension were correlated with a region in the ATL that was more rostral than the region correlated with RTs in the *Shape* dimension. No significant positive correlation was observed. At p < 0.001 uncorrected threshold, additional clusters were identified that are described in the supplementary results.

To illustrate this finding, we examined the functional profile of *Shape-related (*the right posterior ATL region; r-post-ATL, in green on Figure 3) and *Category-related* (the right anterior ATL region; r-ant-ATL, in red on Figure 3) regions. GM measures were extracted from each individual pre-processed structural images using FSL software, and averaged across voxels within each of these 2 clusters, excluding the region of overlap between the two clusters. We ran multiple regressions between each region (r-ant-ATL and r-post-ATL) and *Category* and *Shape* RTs. GM volume in each region was entered as the dependent variable in regression models, and performance in both *Shape* and *Category* tasks were entered as independent variables, together with age, gender, education and total GM volume. R-ant-ATL volume (F6,43=8.1; p<0.001) was significantly predicted by *Category* RT (beta: −0.673, p=0.001) but not by *Shape* RT (beta: −0.020; p=0.927), nor by age, gender, education or total GM volume. R-post-ATL volume (F6,43=4.813; p=0.001) was predicted by *Shape* RT (beta: - 0.598; p=0.016) but not by *Category* RT (beta:-0.072; p=0.740), nor by age, gender, education, or total GM volume. The plots are provided in Figure 3.

### 3.3. GM correlations with RTs in the Same and Different conditions (Table 1, Supplementary Figure 2)

At an FWE-corrected threshold, RTs in the *Same* condition were negatively correlated with GM volume in the right temporal pole, middle temporal gyrus, and inferior temporal gyrus, whereas RTs in the *Different* condition were negatively correlated with the right middle and inferior temporal gyri, i.e. less GM volume was related to slower RTs. There was a large overlap of both clusters (*Same* and *Different*) in the temporal lobe (Supplementary Figure 2). No positive correlation was found with RTs in the *Same* and *Different* conditions.

At p < 0.001 uncorrected threshold, additional negative correlations were found with RTs in the *Same* and *Different* conditions as described in the supplementary results.

### 3.4. WM correlations with RT in Shape, Category, Same, and Different tasks

At an FWE-corrected threshold, RTs in the *Category* dimension were negatively correlated with WM volume in the right temporal lobe (Table 2) i.e. less WM volume was related to slower RTs. This WM region was strictly adjacent to the GM cluster that was correlated negatively with RTs in the *Category* dimension (Figure 4a). To determine what fibers were passing through this region, we explored the anatomical connectivity of the WM-VBM region using tractography-based analyses. No negative correlation was observed with RTs in the *Shape* dimension, as well as the *Same* and *Different* conditions. No significant positive correlation was observed.

**Table 2.**
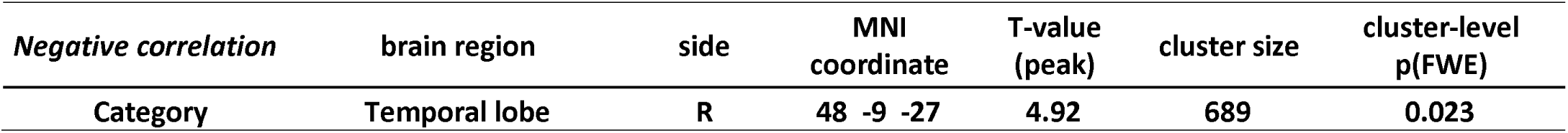
WM correlations with RT in Category at p < 0.05 after FWE correction at the cluster level. Same, Different and Shape conditions were not significant.

| *Negative correlation* | brain region | side | MNI<br>coordinate | T-value<br>(peak) | cluster size | cluster-level<br>p(FWE) |
| --- | --- | --- | --- | --- | --- | --- |
| Category | Temporal lobe | R | 48 -9 -27 | 4.92 | 689 | 0.023 |

**Fig. 4.**
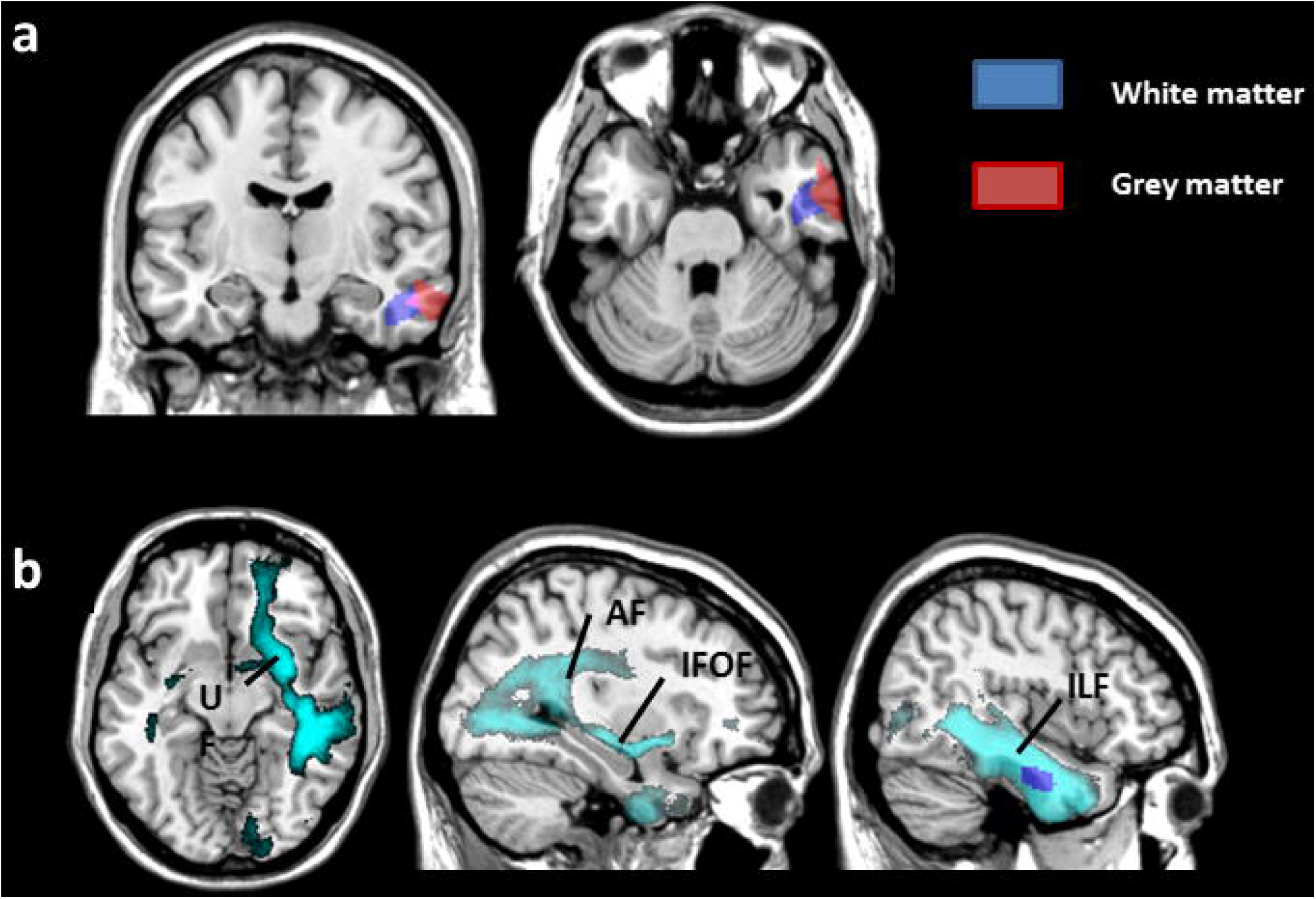
Results from whole-brain WM analysis. p<0.05 after FWE correction. a. Significant regions associated with changes in GM volume (red) and WM volume (blue) related to performance in Category tasks are superimposed on a coronal (left) and axial (right) view. b. The connectome (light blue) represents fibers connecting the right temporal WM region (dark blue) associated with category performance. It includes projection fibers from the right arcuate fasciculus (AF, long segment), inferior fronto-occipital fasciculus (IFOF), uncinate fasciculus (UF), and inferior longitudinal fasciculus (ILF). The axial view is displayed on the left, and the sagittal views are on the right.

At p < 0.001 uncorrected for multiple comparisons, additional negative correlation was found as described in the supplementary results.

### 3.5. Connectivity patterns of the WM-VBM region (Figure 4b)

The connectome representing fibers connecting the right temporal WM region associated with category performance included projection fibers from the right arcuate fasciculus (AF, long segment), inferior fronto-occipital fasciculus (IFOF), uncinate fasciculus (UF), inferior longitudinal fasciculus (ILF), and commissural fibers encompassing the anterior commissure and corpus callosum (splenium).

## 4. Discussion

In this study we performed a voxel-based morphometry study to explore the relationship between semantic and shape categorization tasks and morphometric differences in the brain. Three findings emerge from our work. Firstly, our results revealed a significant correlation between subjects’ performance in terms of RT in all conditions and dimensions, and the volume of the right anterior middle and inferior temporal gyri encompassing the ATL. Secondly, the semantic (*Category*) dimension was associated with a more rostral temporal region than the visuoperceptual (*Shape*) dimension. Finally, WM and connectivity analyses showed a correlation between semantic categorization abilities and WM volume in the right temporal lobe, suggesting the role of the right temporal lobe connections in categorization. Tractography analysis showed that these connections might run through the AF, IFOF, UF, and ILF.

### 4.1. The right anterior middle and inferior temporal gyri and categorization tasks

Interindividual variability in RTs in categorization tasks was related to the GM volume in the right lateral temporal regions. Subjects who were faster to categorize drawings had higher GM volume in the right anterior middle and inferior temporal gyri. To our knowledge, this study is the first to show a correlation between categorization abilities and regional GM volume in healthy participants. This result suggests the role of the lateral part of the right ATL in categorization. Our results are consistent with previous studies that showed a correlation between conceptual processing performances in healthy subjects and resting functional connectivity in the ATL (Wei et al. 2012) in relation to the default mode network.

Previous functional imaging data inconsistently showed the involvement of the ATL during perceptual or semantic categorization tasks. Some authors showed an activation of the ATL (Devlin et al. 2000; Visser et al. 2010a; Binney et al. 2010; Visser et al. 2012), whereas others found an activation of the lateral and/or inferior temporal cortex that was posterior to the ATL (Gerlach et al. 2000; Adams and Janata 2002; Reber et al. 2002; Pernet et al. 2004, 2005; Garcin et al. 2012). The discrepancy of these results may be explained by several factors. First, in fMRI, the observed recruitment of the ATL, a region that is thought to function as a supramodal hub in semantic representation (Patterson et al. 2007), is highly dependent on the contrasting control task that may (Gerlach et al. 2000; Sass et al. 2009) or may not include (Pilgrim et al. 2002; Garcin et al. 2012) a semantic treatment. Second, imaging temporal lobes during classical functional imaging requires a large field of view to ensure whole-brain coverage (Visser et al. 2010b). Finally, evidence of ATL activation is difficult to capture from functional imaging because of susceptibility artifacts caused by variations in magnetic field strength at the interface between brain, bone, and air-filled sinuses; such variations will produce signal loss and distortion (Gorno-Tempini et al. 2002; Visser et al. 2010a). Differences between some of the functional imaging studies and our results may have additional explanations. First, in previous functional imaging studies, the authors examined the regions that were similarly activated across subjects during categorization; they did not explore whether regional activity depends on individual capacities. Second, the correlations found in the present study were based on RTs that, in categorization matching tasks, might correlate with different regions in the temporal lobe than accuracy.

Previous functional imaging data showed the involvement of both the right and left lateral and inferior temporal cortices (Adams and Janata 2002; Pernet et al. 2004; Garcin et al. 2012) and both right and left ATL (Devlin et al. 2000; Visser et al. 2010a; Binney et al. 2010) in categorization tasks. The right lateralization of our findings based on a categorization paradigm using drawings raised the question of a possible hemispheric specialization according to the nature of the stimuli. The possible specialization of the left and right ATLs for verbal versus pictorial semantic representations input is currently under debate in the field (Visser et al. 2010b; Gainotti 2012, 2015). In semantic dementia, left ATL atrophy is correlated with performance in tasks using verbal stimuli (words), whereas right ATL atrophy is correlated with performance in similar tasks using pictorial material (Butler et al. 2009; Acres et al. 2009). Additional anatomic or functional imaging studies of patients with semantic dementia (Butler et al. 2009; Mion et al. 2010; Snowden et al. 2012) and healthy subjects (Thierry et al. 2003; Tsukiura et al. 2006), as well as a recent review on this topic (Gainotti 2015), suggested a verbal/non-verbal dissociation in the ATL. On the contrary, (Pobric et al. 2010) showed that inhibitory repetitive transcranial magnetic stimulation over the right or left temporal pole induces a similar verbal and pictorial (non-verbal) deficit in semantic categorization tasks. A recent meta-analysis on the role of ATL in semantic processing performed by the same group did not find support for lateralization within the ATL but demonstrated that visual object processing often recruits ventral ATL structures, while linguistic and auditory processing recruits lateral ATL structures (Visser et al. 2010b).

Additional studies will be necessary to determine whether there is an actual hemispheric specialization according to the nature of the stimuli and to test whether verbal categorization performances with verbal stimuli are correlated with morphometry of the left ATL.

We cannot exclude that the right lateralization of the main effects in our study can be due to more structural variability on the right ATL than on the left ATL.

Overall, our results complete previous functional imaging findings by demonstrating the relationship between the ability to categorize and the structure of the anterior temporal cortex.

### 4.2. Specialization within the anterior temporal cortex

We showed a rostrocaudal specialization within the temporal lobe: performance in the semantic (*Category*) tasks was associated with more anterior regions of the middle and inferior right temporal gyri than performance in the perceptual (*Shape*) tasks (Figure 3 and supplementary figure 1). These results are in agreement with previous imaging data, suggesting that the posteroventral temporal cortex may encode perceptual categorization, such as categorization based on shape or color, sometimes referred to as “presemantic” (Whatmough et al. 2002), whereas more anterior areas encode semantic categories (Devlin et al. 2005; Binder et al. 2009; Chan et al. 2011; Peelen and Caramazza 2012). However, the shape-related area in our study is more anterior than posteroventral temporal cortex that is usually associated with perceptual categorization. Some authors have proposed that modality specific information is processed in relatively specialized parts of the posterior temporal lobe, whereas the anterior regions are more modality invariant (Visser et al. 2012) or process more abstract/conceptual associations (Bonner and Price 2013). Our findings are consistent with these views, by suggesting a rostrocaudal specialization within the right lateral temporal cortex for processing the *Category* and *Shape* dimensions. Whether this specialization relies on a difference in abstraction between these dimensions, or on the semantic nature of the category task, remains to be tested. Alternatively, the domain-dependent specialization of the anterior versus posterior region of the temporolateral region may support verbal versus non-verbal representations, as participants reported a subvocal verbalization of the semantic category in the *Category* dimension but not in the *Shape* dimension.

### 4.3. Involvement of frontotemporal connections

VBM of the WM and connectivity analyses showed a correlation between RTs in semantic categorization (*Category* dimension) and WM volume in the right temporal lobe. The WM VBM region was adjacent to the GM VBM right temporal region that was correlated with the performance in the *Category* dimension (Figure 4a). The WM VBM region included projection fibers from the right IFOF, UF, long segment of the AF, and ILF. The ILF is associated with object and face recognition, and it is part of the ventral stream (Ortibus et al. 2012; Tavor et al. 2014). Its involvement in our tasks was expected, as subjects had to identify objects to categorize them. According to previous work (Duffau et al. 2005), the IFOF and UF are important pathways for relaying information in semantic memory in the dominant hemisphere. This finding is concordant with a recent morphometry study that found a correlation between the left IFOF and UF and semantic memory performance in healthy subjects (de Zubicaray et al. 2011). Although right-sided, the implication of the IFOF and UF is relevant in the *Category* dimension that relies on the semantic knowledge of the objects to categorize.

IFOF, UF, and AF connect the ATL with the frontal lobe. More specifically, the IFOF and UF connect the ATL with medial and lateral orbitofrontal PFC, whereas the AF connects the ATL with the ventrolateral PFC (Thiebaut de Schotten et al. 2011; Binney et al. 2012; Rojkova et al. 2015). Frontal lobes are most likely involved in categorization tasks, notably in the executive control necessary for categorization. Increasing evidence supported the role of frontal lobes in categorization. Patients with frontal lobe lesions show categorization difficulties (Pribram and Luria 1973; Stuss et al. 1983; Dubois et al. 2000; Fine et al. 2009; Garcin et al. 2012; Lagarde et al. 2015). Functional imaging studies also indicated a role of the lateral PFC for categorization (Tyler et al. 2001; Devlin et al. 2002; Adams and Janata 2002; Vogels et al. 2002; Pilgrim et al. 2002; Reber et al. 2002; Grossman et al. 2002; Pernet et al. 2004; Koenig et al. 2005; Sass et al. 2009; Milton et al. 2009; Garcin et al. 2012; Visser et al. 2012), and electrophysiological recording in primates demonstrated a specific role of the PFC in categorization (Freedman et al. 2003). In agreement with these data, our results of GM volume relationships at an uncorrected threshold showed a positive correlation between RTs in the *Shape* dimension and the right IFG (BA 47), and between RTs in the *Category* dimension and the left inferior and middle frontal gyri (BA 45/46) (see supplementary material). Overall, the correlation of subjects’ performances in the *Category* tasks with a temporal WM region and with the tracts that connect the temporal lobes with the frontal lobe, combined with the correlation of frontal GM volume with categorization tasks, suggest a role of the lateral PFC in these tasks.

## 5. Limitations

We could not exclude that variable processing speed may have influenced our results, because our findings were based on RTs and not accuracy. A previous study performed on 367 healthy subjects found a correlation between processing speed as assessed by the part A of the Trail Making Test (REITAN 1955) and GM volume in the right occipital lobe but no correlation with the temporal GM volume (Ruscheweyh et al. 2013). Studies performed on healthy adults revealed a correlation between processing speed and global WM volume, but no correlation was found with regional WM volume (Penke et al. 2010; Magistro et al. 2015). Finally, our results are concordant with previous studies showing that surgical unilateral resection of the ATL in patients with epilepsy (Lambon Ralph et al. 2012) or inhibition of the ATL induced by repetitive transcranial magnetic stimulation in healthy subjects (Pobric et al. 2010) can increase RTs in semantic assessment tasks. For these reasons, our correlations were unlikely solely caused by processing speed itself.

Additionally, the physiological significance of GM volume correlation remains unclear. For instance, performances negatively correlated with GM volume of the PFC. Correlations between cognition and GM volume, notably in the PFC, do not always respond to the assertion “bigger is better”. Some studies have reported a positive correlation (Yuan and Raz 2014) and others have found a negative correlation (Salat et al. 2002; Goh et al. 2011; Smolker et al. 2015; Aichelburg et al. 2016). The physiological link between cognitive performances and GM volume is not fully understood and may depend on brain maturation and on the synaptic pruning that leads to cortex thinning (Shaw et al. 2006; Dumontheil et al. 2008), as well as on environmental factors, such as training and cognitive stimulation.

## 5. Conclusion

Our results showed the role of the right ATL in categorization abilities in healthy subjects. This study suggested a rostrocaudal specialization in the temporolateral cortex according to the nature of the category. Semantic category judgment was associated with more anterior regions than visuoperceptual category judgment. To our knowledge, this is the first study on the cerebral basis of interindividual variability of categorization abilities. The results add to the current knowledge of the cerebral basis of categorization.

## Acknowledgments

This work was supported by the ‘Fondation pour la Recherche Médicale’ (FRM), [grant numbers FDM20150632801 and DEQ20150331725]. Additional support comes from the ‘Agence Nationale de la Recherche’, [grants numbers ANR-09-RPDOC-004-01 and ANR-13-JSV4-0001-01]. The research leading to these results received funding from the program ‘Investissements d’avenir’ ANR-10-IAIHU-06.

## References

Acres K, Taylor KI, Moss HE, et al(2009) Complementary hemispheric asymmetries in object naming and recognition: a voxel-based correlational study. Neuropsychologia 47:1836–1843. doi: 10.1016/j.neuropsychologia.2009.02.024

Adams RB, Janata P (2002) A comparison of neural circuits underlying auditory and visual object categorization. NeuroImage 16:361–77. doi: 10.1006/nimg.2002.1088

Aichelburg C, Urbanski M, Thiebaut de Schotten M, et al(2016) Morphometry of Left Frontal and Temporal Poles Predicts Analogical Reasoning Abilities. Cereb Cortex N Y N 1991 26:915–932. doi: 10.1093/cercor/bhu254

Anderson AW (2005) Measurement of fiber orientation distributions using high angular resolution diffusion imaging. Magn Reson Med 54:1194–1206. doi: 10.1002/mrm.20667

Andersson JLR, Skare S, Ashburner J (2003) How to correct susceptibility distortions in spin-echo echo-planar images: application to diffusion tensor imaging. NeuroImage 20:870–888. doi: 10.1016/S1053-8119(03)00336-7

Arango-Lasprilla JC, Rivera D, Longoni M, et al(2015) Modified Wisconsin Card Sorting Test (MWCST): Normative data for the Latin American Spanish speaking adult population. NeuroRehabilitation 37:563–590. doi: 10.3233/NRE-151280

Ashburner J (2007) A fast diffeomorphic image registration algorithm. NeuroImage 38:95–113. doi: 10.1016/j.neuroimage.2007.07.007

Ashburner J, Friston KJ (2005) Unified segmentation. NeuroImage 26:839–851. doi: 10.1016/j.neuroimage.2005.02.018

Barsalou LW (1991) Deriving Categories to Achieve Goals. In: Psychology of Learning and Motivation. Elsevier, pp 1–64

Binder JR, Desai RH, Graves WW, Conant LL (2009) Where is the semantic system? A critical review and meta-analysis of 120 functional neuroimaging studies. Cereb Cortex N Y N 1991 19:2767–2796. doi: 10.1093/cercor/bhp055

Binney RJ, Embleton KV, Jefferies E, et al(2010) The ventral and inferolateral aspects of the anterior temporal lobe are crucial in semantic memory: evidence from a novel direct comparison of distortion-corrected fMRI, rTMS, and semantic dementia. Cereb Cortex N Y N 1991 20: 2728–2738. doi: 10.1093/cercor/bhq019

Binney RJ, Parker GJM, Lambon Ralph MA (2012) Convergent connectivity and graded specialization in the rostral human temporal lobe as revealed by diffusion-weighted imaging probabilistic tractography. J Cogn Neurosci 24:1998–2014. doi: 10.1162/jocn_a_00263

Bonner MF, Price AR (2013) Where is the anterior temporal lobe and what does it do? J Neurosci Off J Soc Neurosci 33:4213–4215. doi: 10.1523/JNEUROSCI.0041-13.2013

Butler CR, Brambati SM, Miller BL, Gorno-Tempini M-L (2009) The neural correlates of verbal and nonverbal semantic processing deficits in neurodegenerative disease. Cogn Behav Neurol Off J Soc Behav Cogn Neurol 22:73–80. doi: 10.1097/WNN.0b013e318197925d

Caffarra P, Vezzadini G, Dieci F, et al(2004) Modified Card Sorting Test: normative data. J Clin Exp Neuropsychol 26:246–250. doi: 10.1076/jcen.26.2.246.28087

Callahan BL, Macoir J, Hudon C, et al(2010) Normative data for the pyramids and palm trees test in the Quebec-French population. Arch Clin Neuropsychol Off J Natl Acad Neuropsychol 25:212–217. doi: 10.1093/arclin/acq013

Chan AM, Baker JM, Eskandar E, et al(2011) First-pass selectivity for semantic categories in human anteroventral temporal lobe. J Neurosci Off J Soc Neurosci 31:18119–18129. doi: 10.1523/JNEUROSCI.3122-11.2011

de Zubicaray GI, Rose SE, McMahon KL (2011) The structure and connectivity of semantic memory in the healthy older adult brain. NeuroImage 54:1488–1494. doi: 10.1016/j.neuroimage.2010.08.058

Delis DC, Kaplan E, Kramer JH (2001) D-KEFS. The Psychological Corporation

Dell’acqua F, Scifo P, Rizzo G, et al(2010) A modified damped Richardson-Lucy algorithm to reduce isotropic background effects in spherical deconvolution. NeuroImage 49:1446–1458. doi: 10.1016/j.neuroimage.2009.09.033

Dell’acqua F, Simmons A, Williams SCR, Catani M (2012) Can spherical deconvolution provide more information than fiber orientations? Hindrance modulated orientational anisotropy, a true-tract specific index to characterize white matter diffusion. Hum Brain Mapp. doi: 10.1002/hbm.22080

Devlin JT, Rushworth MFS, Matthews PM (2005) Category-related activation for written words in the posterior fusiform is task specific. Neuropsychologia 43:69–74. doi: 10.1016/j.neuropsychologia.2004.06.013

Devlin JT, Russell RP, Davis MH, et al(2000) Susceptibility-induced loss of signal: comparing PET and fMRI on a semantic task. NeuroImage 11:589–600. doi: 10.1006/nimg.2000.0595

Devlin JT, Russell RP, Davis MH, et al(2002) Is there an anatomical basis for category-specificity? Semantic memory studies in PET and fMRI. Neuropsychologia 40:54–75.

Dubois B, Slachevsky A, Litvan I, Pillon B (2000) The FAB: a Frontal Assessment Battery at bedside. Neurology 55:1621–1626.

Duffau H, Gatignol P, Mandonnet E, et al(2005) New insights into the anatomo-functional connectivity of the semantic system: a study using cortico-subcortical electrostimulations. Brain J Neurol 128:797–810. doi: 10.1093/brain/awh423

Dumontheil I, Burgess PW, Blakemore S-J (2008) Development of rostral prefrontal cortex and cognitive and behavioural disorders. Dev Med Child Neurol 50:168–181. doi: 10.1111/j.1469-8749.2008.02026.x

Fine EM, Delis DC, Dean D, et al(2009) Left frontal lobe contributions to concept formation: a quantitative MRI study of performance on the Delis-Kaplan Executive Function System Sorting Test. J Clin Exp Neuropsychol 31:624–631. doi: 10.1080/13803390802419017

Fischl B, Rajendran N, Busa E, et al(2008) Cortical folding patterns and predicting cytoarchitecture. Cereb Cortex N Y N 1991 18:1973–1980. doi: 10.1093/cercor/bhm225

Folstein MF, Folstein SE, McHugh PR (1975) “Mini-mental state”. A practical method for grading the cognitive state of patients for the clinician. J Psychiatr Res 12:189–198.

Freedman DJ, Riesenhuber M, Poggio T, Miller EK (2003) A comparison of primate prefrontal and inferior temporal cortices during visual categorization. J Neurosci Off J Soc Neurosci 23: 5235–46.

Gainotti G (2012) The format of conceptual representations disrupted in semantic dementia: a position paper. Cortex J Devoted Study Nerv Syst Behav 48:521–529. doi: 10.1016/j.cortex.2011.06.019

Gainotti G (2015) Is the difference between right and left ATLs due to the distinction between general and social cognition or between verbal and non-verbal representations? Neurosci Biobehav Rev 51:296–312. doi: 10.1016/j.neubiorev.2015.02.004

Gamboz N, Coluccia E, Iavarone A, Brandimonte MA (2009) Normative data for the Pyramids and Palm Trees Test in the elderly Italian population. Neurol Sci Off J Ital Neurol Soc Ital Soc Clin Neurophysiol 30:453–458. doi: 10.1007/s10072-009-0130-y

Garcin B, Volle E, Dubois B, Lévy R (2012) Similar or different ⍰? The role of the ventrolateral prefrontal cortex in similarity detection. PloS One

Gerlach C, Law I, Gade A, Paulson OB (2000) Categorization and category effects in normal object recognition: a PET study. Neuropsychologia 38:1693–703.

Goh S, Bansal R, Xu D, et al(2011) Neuroanatomical correlates of intellectual ability across the life span. Dev Cogn Neurosci 1:305–312. doi: 10.1016/j.dcn.2011.03.001

Gorno-Tempini ML, Hutton C, Josephs O, et al(2002) Echo time dependence of BOLD contrast and susceptibility artifacts. NeuroImage 15:136–142. doi: 10.1006/nimg.2001.0967

Grossman M, Smith EE, Koenig P, et al(2002) The neural basis for categorization in semantic memory. NeuroImage 17:1549–1561.

Harrison AG, Armstrong IT, Harrison LE, et al(2014) Comparing Canadian and American normative scores on the Wechsler Adult Intelligence Scale-Fourth Edition. Arch Clin Neuropsychol Off J Natl Acad Neuropsychol 29:737–746. doi: 10.1093/arclin/acu048

Homack S, Lee D, Riccio CA (2005) Test review: Delis-Kaplan executive function system. J Clin Exp Neuropsychol 27:599–609. doi: 10.1080/13803390490918444

Howard D, Patterson K (1992) Pyramids and palm tress: A test of semantic access from pictures and words.

Klein A, Andersson J, Ardekani BA, et al(2009) Evaluation of 14 nonlinear deformation algorithms applied to human brain MRI registration. NeuroImage 46:786–802. doi: 10.1016/j.neuroimage.2008.12.037

Klein LA, Buchanan JA (2009) Psychometric properties of the Pyramids and Palm Trees Test. J Clin Exp Neuropsychol 31:803–808. doi: 10.1080/13803390802508926

Koenig P, Smith EE, Glosser G, et al(2005) The neural basis for novel semantic categorization. NeuroImage 24:369–383. doi: 10.1016/j.neuroimage.2004.08.045

Lagarde J, Valabrègue R, Corvol J-C, et al(2015) Why do patients with neurodegenerative frontal syndrome fail to answer: “In what way are an orange and a banana alike?” Brain J Neurol 138:456–471. doi: 10.1093/brain/awu359

Lambon Ralph MA, Ehsan S, Baker GA, Rogers TT (2012) Semantic memory is impaired in patients with unilateral anterior temporal lobe resection for temporal lobe epilepsy. Brain J Neurol 135:242–258. doi: 10.1093/brain/awr325

Leemans A, Jones DK (2009) The B-matrix must be rotated when correcting for subject motion in DTI data. Magn Reson Med 61:1336–1349. doi: 10.1002/mrm.21890

Luo H, Husain FT, Horwitz B, Poeppel D (2005) Discrimination and categorization of speech and non-speech sounds in an MEG delayed-match-to-sample study. NeuroImage 28:59–71. doi: 10.1016/j.neuroimage.2005.05.040

Magistro D, Takeuchi H, Nejad KK, et al(2015) The Relationship between Processing Speed and Regional White Matter Volume in Healthy Young People. PloS One 10:e0136386. doi: 10.1371/journal.pone.0136386

Mattioli F, Stampatori C, Bellomi F, et al(2014) Assessing executive function with the D-KEFS sorting test: normative data for a sample of the Italian adult population. Neurol Sci Off J Ital Neurol Soc Ital Soc Clin Neurophysiol 35:1895–1902. doi: 10.1007/s10072-014-1857-7

Mazziotta JC, Toga AW, Evans A, et al(1995) A probabilistic atlas of the human brain: theory and rationale for its development. The International Consortium for Brain Mapping (ICBM). NeuroImage 2:89–101.

Merck C, Charnallet A, Auriacombe S, et al(2011) La batterie d’évaluation des connaissances sémantiques du GRECO (BECS-GRECO)M: validation et données normatives. Rev Neuropsychol 3:235–255. doi: 10.3917/rne.034.0235

Milner B (1963) Effect of Different Brain Lesions on Card Sorting. Arch. Neurol. 100–110.

Milton F, Wills AJ, Hodgson TL (2009) The neural basis of overall similarity and single-dimension sorting. NeuroImage 46:319–326. doi: 10.1016/j.neuroimage.2009.01.043

Mion M, Patterson K, Acosta-Cabronero J, et al(2010) What the left and right anterior fusiform gyri tell us about semantic memory. Brain J Neurol 133:3256–3268. doi: 10.1093/brain/awq272

Ortibus E, Verhoeven J, Sunaert S, et al(2012) Integrity of the inferior longitudinal fasciculus and impaired object recognition in children: a diffusion tensor imaging study. Dev Med Child Neurol 54:38–43. doi: 10.1111/j.1469-8749.2011.04147.x

Patterson K, Nestor PJ, Rogers TT (2007) Where do you know what you know? The representation of semantic knowledge in the human brain. Nat Rev Neurosci 8:976–987. doi: 10.1038/nrn2277

Peelen MV, Caramazza A (2012) Conceptual Object Representations in Human Anterior Temporal Cortex. J Neurosci 32:15728–15736. doi: 10.1523/JNEUROSCI.1953-12.2012

Penke L, Muñoz Maniega S, Murray C, et al(2010) A general factor of brain white matter integrity predicts information processing speed in healthy older people. J Neurosci Off J Soc Neurosci 30:7569–7574. doi: 10.1523/JNEUROSCI.1553-10.2010

Pernet C, Celsis P, Démonet J-F (2005) Selective response to letter categorization within the left fusiform gyrus. NeuroImage 28:738–44. doi: 10.1016/j.neuroimage.2005.06.046

Pernet C, Franceries X, Basan S, et al(2004) Anatomy and time course of discrimination and categorization processes in vision: an fMRI study. NeuroImage 22:1563–77. doi: 10.1016/j.neuroimage.2004.03.044

Pilgrim LK, Fadili J, Fletcher P, Tyler LK (2002) Overcoming confounds of stimulus blocking: an event-related fMRI design of semantic processing. NeuroImage 16:713–723.

Pobric G, Jefferies E, Ralph MAL (2010) Amodal semantic representations depend on both anterior temporal lobes: evidence from repetitive transcranial magnetic stimulation. Neuropsychologia 48:1336–1342. doi: 10.1016/j.neuropsychologia.2009.12.036

Pribram KH, Luria AR (1973) Psychophysiology of the Frontal Lobes. Academic P

Rami L, Serradell M, Bosch B, et al(2008) Normative data for the Boston Naming Test and the Pyramids and Palm Trees Test in the elderly Spanish population. J Clin Exp Neuropsychol 30:1–6. doi: 10.1080/13803390701743954

Reber PJ, Wong EC, Buxton RB (2002) Comparing the brain areas supporting nondeclarative categorization and recognition memory. Brain Res Cogn Brain Res 14:245–57.

Reitan RM (1955) The relation of the trail making test to organic brain damage. J Consult Psychol 19:393–394.

Rojkova K, Volle E, Urbanski M, et al(2015) Atlasing the frontal lobe connections and their variability due to age and education: a spherical deconvolution tractography study. Brain Struct Funct. doi: 10.1007/s00429-015-1001-3

Ruscheweyh R, Deppe M, Lohmann H, et al(2013) Executive performance is related to regional gray matter volume in healthy older individuals. Hum Brain Mapp 34:3333–3346. doi: 10.1002/hbm.22146

Salat DH, Kaye JA, Janowsky JS (2002) Greater orbital prefrontal volume selectively predicts worse working memory performance in older adults. Cereb Cortex N Y N 1991 12:494–505.

Sass K, Sachs O, Krach S, Kircher T (2009) Taxonomic and thematic categories: Neural correlates of categorization in an auditory-to-visual priming task using fMRI. Brain Res 1270: 78–87. doi: 10.1016/j.brainres.2009.03.013

Schmahmann JD, Pandya DN, Wang R, et al(2007) Association fibre pathways of the brain: parallel observations from diffusion spectrum imaging and autoradiography. Brain J Neurol 130: 630–653. doi: 10.1093/brain/awl359

Shan I-K, Chen Y-S, Lee Y-C, Su T-P (2008) Adult normative data of the Wisconsin Card Sorting Test in Taiwan. J Chin Med Assoc JCMA 71:517–522. doi: 10.1016/S1726-4901(08)70160-6

Shaw P, Greenstein D, Lerch J, et al(2006) Intellectual ability and cortical development in children and adolescents. Nature 440:676–679. doi: 10.1038/nature04513

Sigala N, Logothetis NK (2002) Visual categorization shapes feature selectivity in the primate temporal cortex. Nature 415:318–20. doi: 10.1038/415318a

Smith SM, Jenkinson M, Woolrich MW, et al(2004) Advances in functional and structural MR image analysis and implementation as FSL. NeuroImage 23 Suppl 1: S208−219. doi: 10.1016/j.neuroimage.2004.07.051

Smolker HR, Depue BE, Reineberg AE, et al(2015) Individual differences in regional prefrontal gray matter morphometry and fractional anisotropy are associated with different constructs of executive function. Brain Struct Funct 220:1291–1306. doi: 10.1007/s00429-014-0723-y

Snowden JS, Thompson JC, Neary D (2012) Famous people knowledge and the right and left temporal lobes. Behav Neurol 25:35–44. doi: 10.3233/BEN-2012-0347

Stuss DT, Benson DF, Kaplan EF, et al(1983) The involvement of orbitofrontal cerebrum in cognitive tasks. Neuropsychologia 21:235–248.

Tavor I, Yablonski M, Mezer A, et al(2014) Separate parts of occipito-temporal white matter fibers are associated with recognition of faces and places. NeuroImage 86:123–130. doi: 10.1016/j.neuroimage.2013.07.085

Thiebaut de Schotten M, Ffytche DH, Bizzi A, et al(2011) Atlasing location, asymmetry and inter-subject variability of white matter tracts in the human brain with MR diffusion tractography. NeuroImage 54:49–59. doi: 10.1016/j.neuroimage.2010.07.055

Thierry G, Giraud AL, Price C (2003) Hemispheric dissociation in access to the human semantic system. Neuron 38:499–506.

Tournier J-D, Calamante F, Gadian DG, Connelly A (2004) Direct estimation of the fiber orientation density function from diffusion-weighted MRI data using spherical deconvolution. NeuroImage 23:1176–1185. doi: 10.1016/j.neuroimage.2004.07.037

Tsukiura T, Mochizuki-Kawai H, Fujii T (2006) Dissociable roles of the bilateral anterior temporal lobe in face-name associations: an event-related fMRI study. NeuroImage 30: 617–626. doi: 10.1016/j.neuroimage.2005.09.043

Tyler LK, Russell R, Fadili J, Moss HE (2001) The neural representation of nouns and verbs: PET studies. Brain J Neurol 124:1619–34.

Uylings HBM, Rajkowska G, Sanz-Arigita E, et al(2005) Consequences of large interindividual variability for human brain atlases: converging macroscopical imaging and microscopical neuroanatomy. Anat Embryol (Berl) 210:423–431. doi: 10.1007/s00429-005-0042-4

Visser M, Embleton KV, Jefferies E, et al(2010a) The inferior, anterior temporal lobes and semantic memory clarified: novel evidence from distortion-corrected fMRI. Neuropsychologia 48:1689–1696. doi: 10.1016/j.neuropsychologia.2010.02.016

Visser M, Jefferies E, Embleton KV, Lambon Ralph MA (2012) Both the middle temporal gyrus and the ventral anterior temporal area are crucial for multimodal semantic processing: distortion-corrected fMRI evidence for a double gradient of information convergence in the temporal lobes. J Cogn Neurosci 24:1766–1778. doi: 10.1162/jocn_a_00244

Visser M, Jefferies E, Lambon Ralph MA (2010b) Semantic processing in the anterior temporal lobes: a meta-analysis of the functional neuroimaging literature. J Cogn Neurosci 22:1083–1094. doi: 10.1162/jocn.2009.21309

Vogels R, Sary G, Dupont P, Orban GA (2002) Human brain regions involved in visual categorization. NeuroImage 16:401–14. doi: 10.1006/nimg.2002.1109

Wechsler D (2008) Wechsler Adult Intelligence Scale - Fourth Edition (WAIS - IV) Administration and Scoring Manual., Pearson. San Antonio

Wei T, Liang X, He Y, et al(2012) Predicting conceptual processing capacity from spontaneous neuronal activity of the left middle temporal gyrus. J Neurosci Off J Soc Neurosci 32:481–489. doi: 10.1523/JNEUROSCI.1953-11.2012

Whatmough C, Chertkow H, Murtha S, Hanratty K (2002) Dissociable brain regions process object meaning and object structure during picture naming. Neuropsychologia 40:174–186.

Wisdom NM, Mignogna J, Collins RL (2012) Variability in Wechsler Adult Intelligence Scale-IV subtest performance across age. Arch Clin Neuropsychol Off J Natl Acad Neuropsychol 27:389–397. doi: 10.1093/arclin/acs041

Yuan P, Raz N (2014) Prefrontal cortex and executive functions in healthy adults: a meta-analysis of structural neuroimaging studies. Neurosci Biobehav Rev 42:180–192. doi: 10.1016/j.neubiorev.2014.02.005

